# SELECTIVELY MANIPULATING SOFTNESS PERCEPTION OF MATERIALS THROUGH SOUND SYMBOLISM

**DOI:** 10.1101/2023.10.18.562669

**Authors:** H. Nalbantoğlu, B. M. Hazır, D. N. Dövencioğlu

## Abstract

Cross-modal interactions between auditory and haptic perception manifest themselves in language, such as sound symbolic words: crunch, splash, and creak. Several studies have shown strong associations between sound symbolic words, shapes (e.g., Bouba/Kiki effect), and materials. Here, we identified these material associations in Turkish sound symbolic words and then tested for their effect on softness perception. First, we used a rating task in a semantic differentiation method to extract the perceived softness dimensions from words and materials. We then tested whether Turkish onomatopoeic words can be used to manipulate the perceived softness of everyday materials such as honey, silk, or sand across different dimensions of softness. In the first preliminary study, we used 40 material videos and 29 adjectives in a rating task with a semantic differentiation method to extract the main softness dimensions. A principal component analysis revealed 7 softness components, including Deformability, Viscosity, Surface Softness, and Granularity, in line with the literature. The second preliminary study used 47 Turkish onomatopoeic words and 31 adjectives in the same rating task. Again, the findings aligned with the literature, revealing dimensions such as Fluidity, Granularity, and Surface Softness. However, no factors related to Deformability were found due to the absence of sound symbolic words in this category. Next, we paired the onomatopoeic words and material videos based on their associations with each softness dimension. We conducted a new rating task, synchronously presenting material videos and spoken onomatopoeic words. We hypothesized that congruent word-video pairs would produce significantly higher ratings for dimension-related adjectives, while incongruent word-video pairs would decrease these ratings, and the ratings of unrelated adjectives would remain the same. Our results revealed that onomatopoeic words selectively alter the perceived material qualities, providing evidence and insight into the cross-modality of perceived softness.

## 1 Introduction

Arbitrariness in language proposes a lack of any inherent connection between the sounds of words and their meanings (Imai et al., 2014); however, sound symbolism challenges this notion by asserting that there exists a non-arbitrary relationship between sounds and meaning for some words. A subset of these sound-symbolic words consists of onomatopoeia, words that sound just like the things they refer to. The onomatopoeic words are frequently used in sound-symbolism research since, contrary to the general belief, they are informative about the sounds they refer to and provide cross-modal information about the objects. For instance, the word "crispy" not only conveys the auditory characteristics of the sound produced by crispy objects but also suggests certain tactile and potentially visual properties associated with these objects.

One of the classical examples of these non-arbitrary sound symbolic relationships concerning shape perception is called the "Bouba-Kiki" effect (Köhler, 1929; Maurer et al., 2006; Ozturk et al., 2013; Peiffer-Smadja & Cohen, 2019). In the earlier findings, participants often associated angular objects with pseudowords “Kiki” and round objects with “Bouba” (Ramachandran & Hubbard, 2001). Since then, the effect has been demonstrated across different cultural backgrounds and language groups, implying the universality of sound symbolism (e.g., Bremner et al., 2013; Ćwiek et al., 2021).

In addition to these, Etzi et al. (2016) found that round-shaped sounds (as in “Bouba”) were more related to smoother textures, while sharp-transient sounds (as in “Kiki”) were more related to rougher textures. There is also evidence for strong associations between sounds and material perception in relatively recent research (Jousmäki and Hari, 1998; Watanabe et al., 2012; Sakamoto & Watanabe, 2018; Wong et al., 2022). For instance, Sakamoto and Watanabe (2018) found specific relationships between sounds and tactile ratings in the Japanese language, such as /p/, /b/, and /n/ consonants being more related to soft materials while /ts/ and /k/ being more related to hard materials. When they asked participants to generate sound-symbolic words in Japanese (including novel pseudowords) while touching a variety of materials, the results revealed that voiced consonants (e.g., /dz/ and /g/) were more likely to be associated with roughness and voiceless consonants (e.g., /ʦ/, and /s/) with smoothness. Further evidence supporting the crossmodal interactions between haptic and auditory signals comes from the Parchment-skin illusion (Jousmäki & Hari, 1998; Guest et al., 2002; Fujisaki, 2015), where an auditory signal’s frequency and sound levels alter tactile roughness perception. For instance, researchers simultaneously presented sounds of varying frequencies and levels while participants rubbed their hands; they found that higher frequencies and sound levels led to the perception of more paper-like and rougher skin (Guest et al., 2002).

In another study, Lo et al. (2017) conducted two experiments investigating the relationship of fricative-plosive consonants to spiky-curved shapes and to rough-smooth tactile surfaces. They found no relationship between the fricative and plosive sounds and spiky-curved shapes. However, their second experiment results revealed that participants were more likely to associate fricative-consonant speech sounds (i.e., /f/, /h/, /s/) with rough-textured materials and plosive-consonant speech sounds (i.e., /b/, /p/, /t/) with smooth-textured materials. The relationship between speech sounds and food texture was also investigated in the literature. For example, Hanada (2019) used Japanese onomatopoeic words to extract 15 perceptual dimensions of food texture, such as smoothness, adhesiveness, or wateriness. In a later study, Hanada (2023) investigated the fabrics’ haptic dimensions and feelings of luxuriousness and pleasantness using onomatopoeia in Japanese. In terms of perceived fabric luxuriousness and pleasantness, the likelihood of having the /s/, /h/, /m/, and /u/ sounds were found to increase, while the /k/, /g/, /z/, /p/, and /a/ sounds were found to decrease. They also reported that /k/, /g/, /z/, /p/, and /a/ sounds were more associated with the cheapness and unpleasantness of fabrics.

The studies above provide evidence of sound symbolism’s role in tactile perception. However, the characteristics of touch signals are not limited to the tactile information from two-dimensional textures: Haptic, visual, and auditory research shows that human softness perception has multiple dimensions from viscous, granular, deformable materials to materials with soft surfaces (Cavdan et al., 2019; 2021; Dovencioğlu et al., 2022). These softness dimensions are also closely related to how humans interact with materials and surfaces in their environment using exploratory procedures, which refer to a set of hand movements used to explore various material properties (Lederman & Klatzky, 1987, Dovencioğlu et al., 2022). Some examples include using pressing deformable materials, rotating granular materials, and stroking materials with soft surfaces.

These findings might suggest a connection between sound symbolism and the multidimensionality of perceived softness, especially concerning materials that go beyond textiles (e.g., Hanada, 2023). Studies that relate sound symbolism to material perception are mostly limited to the Japanese language, which has a diverse vocabulary of onomatopoeic words relating to tactile perception (Sakamoto & Watanabe, 2018; Watanabe et al., 2012; Wong et al., 2022). Turkish is also rich in the number of onomatopoeic words that are used colloquially. Many of these onomatopoeic words describe the softness-related properties of materials, such as “pıtır pıtır” (with a patter) for granular, “tiril tiril” (floaty) for silky, and “vıcık vıcık” (slushy) for gooey materials. Moreover, these words may even have subtle non-arbitrary variations, depending on the nature of the action or movement, such as “şarıl şarıl” for water that flows abundantly with splashing sounds, ’şırıl şırıl’ for water that flows pleasantly in a small amount but continuously, and ’şorul şorul,’ although a less common onomatopoeia, for water that flows abundantly and loudly (Zulfikar, 1995). Despite a significant number of examples, there is no research looking for systematic associations between softness perception and onomatopoeia in Turkish.

In the current study, we aim to explore the potential influence of Turkish onomatopoeic words on the perception of softness in everyday materials. To this end, we created congruent and incongruent pairs of videos of everyday materials and spoken Turkish onomatopoeic words for each softness dimension. We conducted two preliminary studies using the semantic differentiation method with 1) material videos and 2) onomatopoeic words. We extracted perceived softness factors from these studies and determined the congruency of onomatopoeic words and material videos based on their ratings for softness-related adjectives. Next, we used these pairs in Experiments 1 and 2. We hypothesized that pairing materials with onomatopoeic words would selectively alter their perceived material qualities, i.e., congruent pairs with high-rated onomatopoeic words would increase the mean ratings of adjectives related to the softness dimensions of materials, whereas incongruent pairs with low-rated onomatopoeic words would decrease the mean ratings.

## 2 Materials and Methods

We conducted two preliminary studies (Supplementary Study 1 and 2) to select the material videos and onomatopoeic words that were used in Experiments 1 and 2. We used the semantic differentiation method in these preliminary studies with 29 and 21 softness-related adjectives, respectively. First, participants rated 40 material videos in an online study (Supplementary Study 1). Next, a different group of participants rated 27 onomatopoeic words in the lab (Supplementary Study 2). Principal Component Analyses revealed 7 softness-related factors in the first and 4 softness-related factors in the second study. Stimulus selection based on these results is described below.

### 2.1 Ethics Statement

The research was approved by the METU Human Subjects Ethics Committee and in accordance with the Declaration of Helsinki.

### 2.2 Experiment 1

Experiment 1 is conducted to determine the baseline judgments for the materials with a smaller set of adjectives. These judgments are then put to the test with congruent and incongruent onomatopoeic words in Experiment 2.

#### 2.2.1 Participants

We recruited 23 participants (M = 21.13, SD = 1.42, 2 Males) for the experiment via the SONA research sign-up system. Participants gave informed consent before the experiment. After completion, they were compensated for their time with course credits. All participants had normal or corrected to normal vision. None of the participants reported hearing loss or any other auditory condition.

#### 2.2.2 Stimulus Selection: Common Dimensions and Adjectives

The selection of adjectives for this experiment was informed by the common factors extracted from Supplementary Study 1 and Supplementary Study 2. The four common factors were Viscosity, Surface Softness, Granularity, and Roughness (Table 1, first column). We selected 13 adjectives that loaded to the same factor in both studies (Table 1, second column). Specifically, we selected 6 adjectives for Viscosity (gelatinous, slimy, sticky, gooey, slippery, moisturous), three adjectives for Surface Softness (silky, velvety, hairy), three adjectives for Granularity (sandy, powdery, granular), and one adjective for Roughness (roughened) dimensions.

**Table 1.**
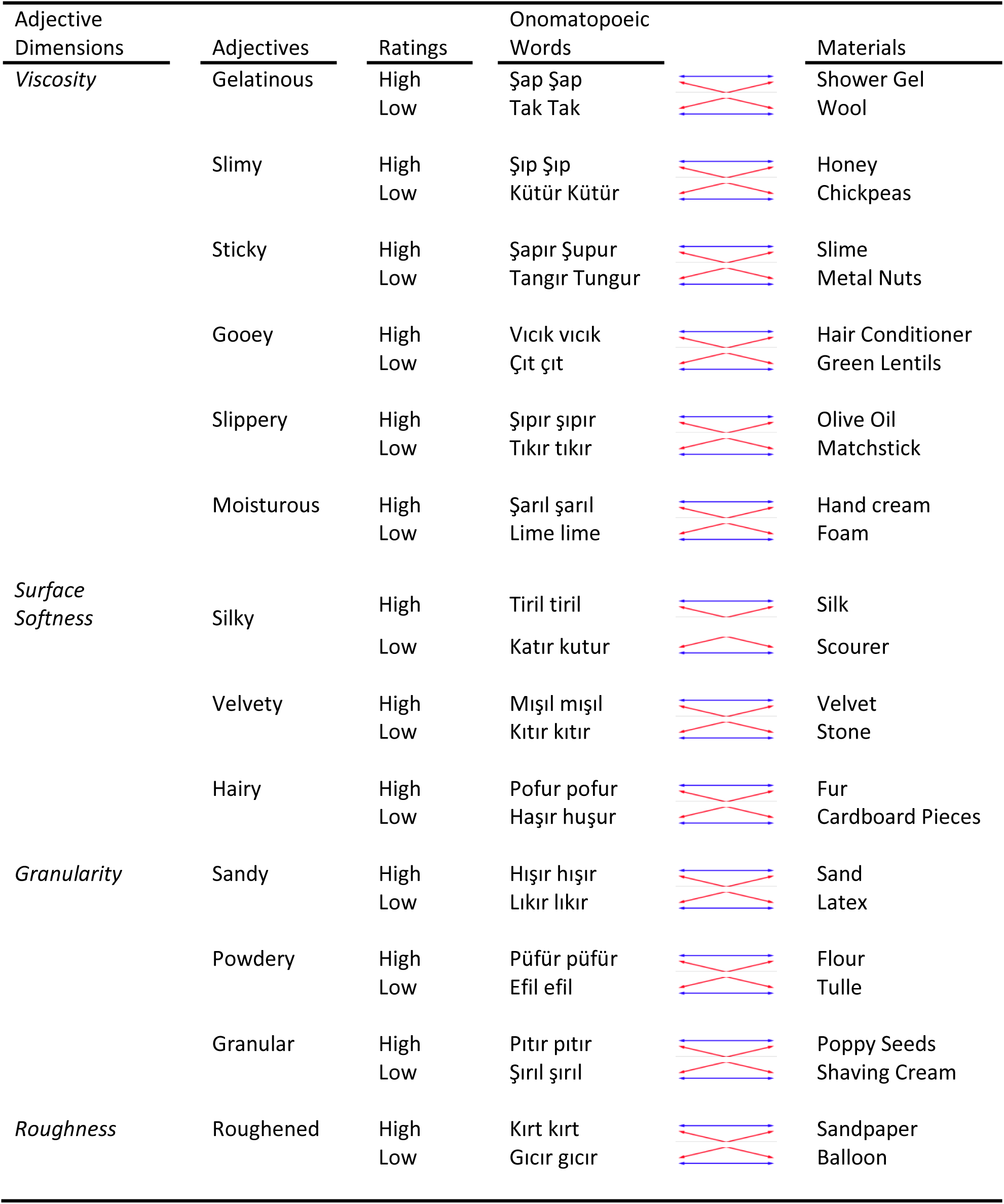
Adjectives, Materials, and Onomatopoeic Words Selected for Each Dimension. High-and low-rated onomatopoeic words and materials are listed for the corresponding adjectives. Blue arrows indicate congruent pairs (both high-or low-rated words and materials), whereas red arrows mark incongruent pairs (e.g., a low-rated word with a high-rated material).

From this adjective list, we continued to select the material videos. For each of the 13 adjectives, two material videos (one rated high and one rated low) from Study 1 were selected. A high rating was defined as a mean rating above 3.5 (out of 7) for a specific adjective. For instance, the shower gel video received a high rating for gelatinous with a mean score of 6.52 (SEM = 0.27). Conversely, a low rating was defined as a mean rating below 3.5. For example, the material video for wool received a low rating for gelatinous with a mean score of 1.35 (SEM = 0.12). The complete list of materials selected for the 13 adjectives can be found in **Table 1**. The resulting material videos also represented the same four dimensions with the adjectives. Overall, we selected 12 material videos for Viscosity, 6 material videos each for Surface Softness and Granularity, and 2 material videos for Roughness dimensions. The adjective scaly was not used for the Granularity dimension since all material ratings for scaly were lower than 3.5 (with a highest of 2.9), resulting in no high-rated materials.

#### 2.2.3 Procedure

The experiment was coded in MATLAB R2020b using Psychtoolbox and consisted of 26 material videos presented in a loop synchronously with the 13 adjectives one by one. One experimental block consisted of 338 trials for each participant. The experiment was conducted using an HP Pavilion TPC-F123-MT computer and HP 2011x 20-inch LED Monitor.

Participants were instructed to rate the adjectives they saw on the screen based on the material videos. A 7-point Likert scale was used for the ratings (1 = "*Not at all”*, 7 = “Very”). A sample trial from Experiment 1 is illustrated in Figure 1.

**Figure 1.**
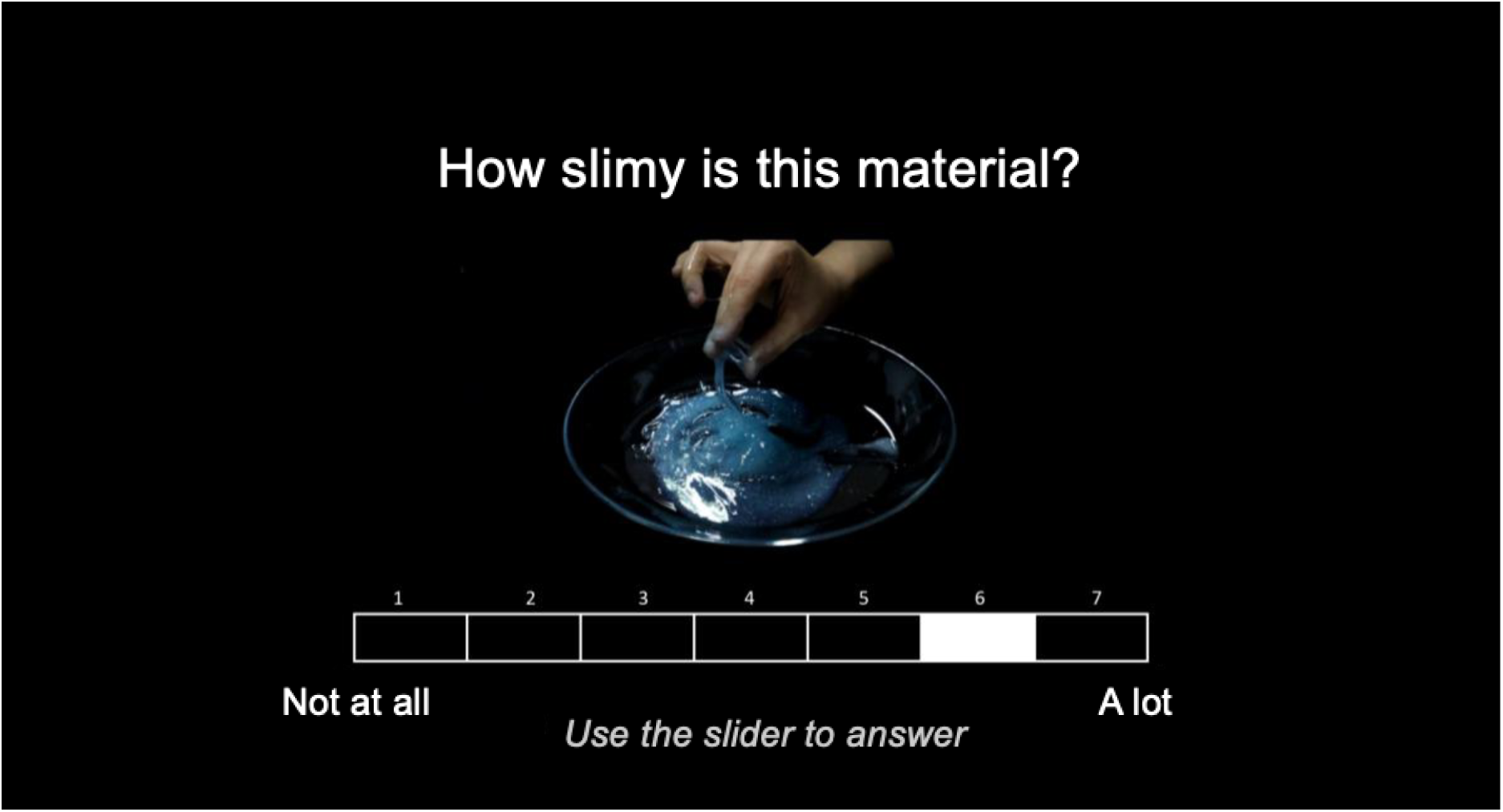
Screenshot from a sample trial in Experiment 1. While the material video of the “shower gel” is playing on the screen, participants are asked to rate the adjective “slimy (sümüksü)” from 1 to 7 based on the material.

#### 2.2.4 Results

For each of the 13 adjectives, we calculated the mean ratings of the four material dimensions (Viscosity, Granularity, Surface Softness, and Roughness) and used them as baseline measurements for the mean ratings in Experiment 2 (Table 2 and Supplementary Figure 5, red lines). We averaged the ratings across within-dimension adjectives to come up with the adjective dimensions (Table 3, rows). Similarly, average ratings across the within-dimension materials gave us the material dimensions (Table 3, columns). All Adjective Dimensions, except for the Roughness Adjective, received the highest mean ratings from their own material dimensions (darker cells in Table 3). Specifically, Viscosity Adjectives (*M* = 3.72, *SE* = 0.06) received the highest mean rating from the Viscosity Materials, Surface Softness Adjectives (*M* = 3.24, *SE* = 0.12) received the highest mean rating from Surface Softness Materials, and Granularity Adjectives (*M* = 3.83, *SE* = 0.14) had the highest mean rating for the Granularity Materials. However, Roughness Adjectives had the highest mean rating for the Surface Softness Materials (*M* = 4, *SE* = 0.17) and the second highest for Roughness Materials (*M* = 3.7, *SE* = 0.33).

**Table 2.**
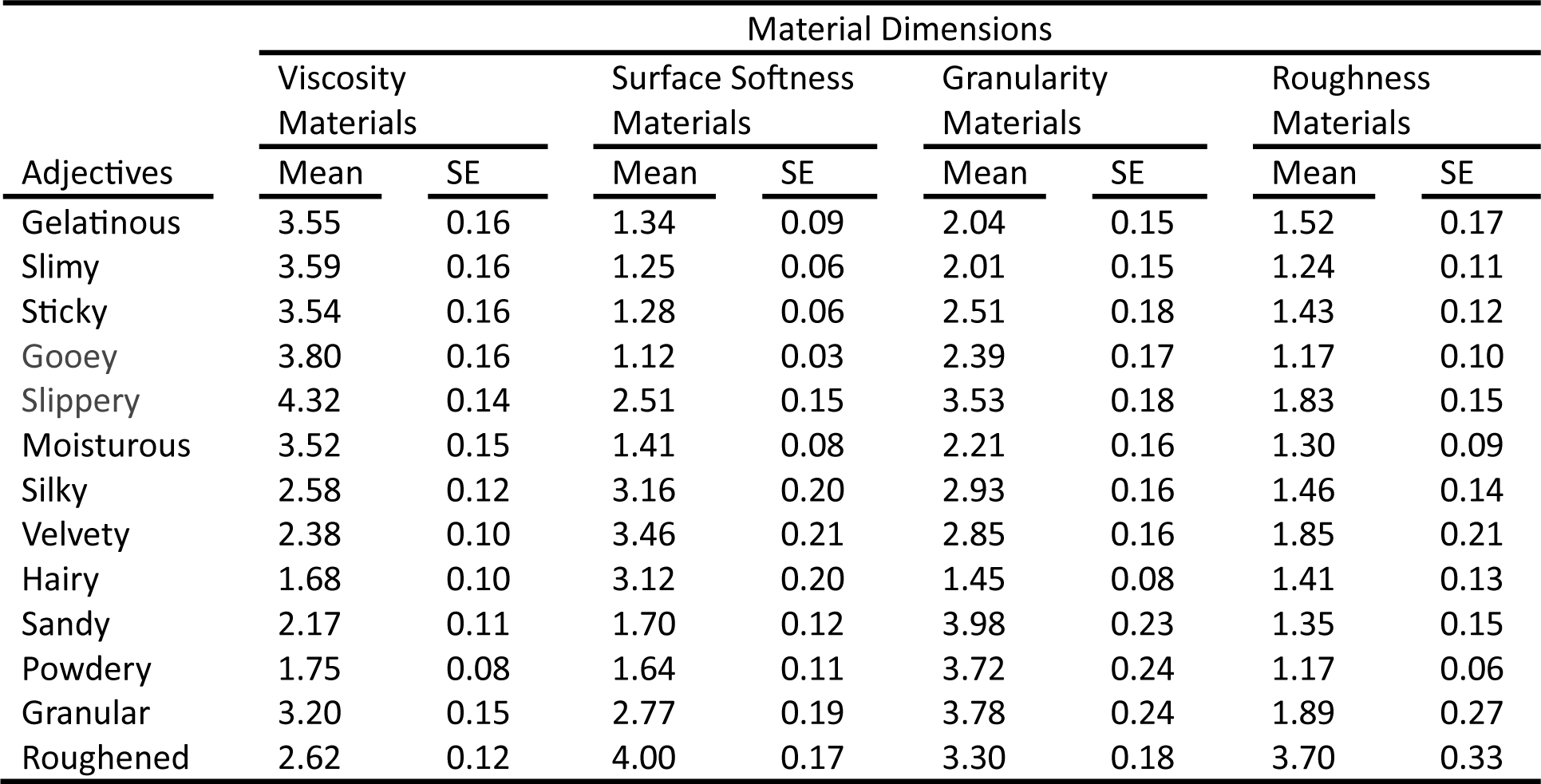
Mean Ratings and Standard Errors of Adjectives Across Material Dimensions.

**Table 3.** Mean Ratings and Standard Errors of Adjective Dimensions Across Material Dimensions.

| Adjective Dimensions | Material Dimensions |  |  |  |  |  |  |  |
| --- | --- | --- | --- | --- | --- | --- | --- | --- |
|  | Viscosity Materials |  | Surface Softness Materials |  | Granularity Materials |  | Roughness Materials |  |
|  | Mean | SE | Mean | SE | Mean | SE | Mean | SE |
| Viscosity Adjectives | 3.72 | 0.06 | 1.48 | 0.04 | 2.45 | 0.07 | 1.42 | 0.05 |
| Surface Softness Adjectives | 2.21 | 0.06 | 3.24 | 0.12 | 2.41 | 0.09 | 1.57 | 0.09 |
| Granularity Adjectives | 2.37 | 0.07 | 2.04 | 0.09 | 3.83 | 0.14 | 1.47 | 0.11 |
| Roughness Adjectives | 2.62 | 0.12 | 4.00 | 0.17 | 3.30 | 0.18 | 3.70 | 0.33 |

When we look at the individual adjectives, we observe the same pattern of higher values for within-dimension ratings. All Viscosity Adjectives received the highest mean ratings from the Viscosity Materials (Table 2, leftmost column, first 6 adjectives). These adjectives were gelatinous (*M* = 3.55, *SE* = 0.16), slimy (*M* = 3.59, *SE* = 0.16), sticky (*M* = 3.54, *SE* = 0.16), gooey (*M* = 3.8, *SE* = 0.16), slippery (*M* = 4.32, *SE* = 0.14), and moisturous (*M* = 3.52, *SE* = 0.15). Similarly, silky (*M* = 3.16, *SE* = 0.2) and velvety (*M* = 3.46, *SE* = 0.21) had the highest mean ratings for the Surface Softness Materials, while sandy (*M* = 3.98, *SE* = 0.23) and powdery (*M* = 3.72, *SE* = 0.24) had the highest mean ratings for the Granularity Materials. Lastly, the adjective roughened, which is the only roughness adjective, received the highest mean rating (*M* = 4, *SE* = 0.17) from the Surface Softness Materials, but it received the second highest mean rating (*M* = 3.7, *SE* = 0.33) from the Roughness Materials.

Overall, the highest mean rating was observed for the adjective slippery (*M* = 4.32, *SE* = 0.14) for the Viscosity Materials, whereas the adjective gooey (*M* = 1.12, *SE* = 0.03) received the lowest mean rating with the Surface Softness Materials.

### 2.3 Experiment 2

#### 2.3.1 Participants

30 participants (*M* = 22.5, *SD* = 2.52, 11 Males) were recruited through the Middle East Technical University research sign-up system. Participants gave written informed consent before the experiment. After completion, they were compensated for their time with course credits. Only one participant was left-handed, and all participants had normal or corrected to normal vision. None of the participants reported hearing loss or any other auditory condition.

#### 2.3.2 Stimulus Selection: Onomatopoeic Words

The adjectives and the material videos used in this experiment were the same as those used in Experiment 1. Additionally, we included 26 onomatopoeic words in this experiment. Similar to the selection process of the material videos in Experiment 1, two onomatopoeic words (one high-rated and one low-rated) were selected for each of the 13 adjectives based on the findings of Supplementary Study 2. A high rating was defined as a mean rating above 3.5 (out of 7) for a specific adjective. For instance, the onomatopoeic word “şap şap” received a high rating for gelatinous with a mean score of 4.7 (SEM = 0.36). Conversely, a low rating was defined as a mean rating below 3.5. For example, “tak tak” received a low rating for gelatinous with a mean score of 1.1 (SEM = 0.1). The complete list of onomatopoeic words selected for the 13 adjectives can be found in Table 1.

The material videos and onomatopoeic words were matched congruently (both are rated high, or both are rated low for a given adjective) or incongruently (one rated high when the other is rated low for a given adjective). We ended up with 4 material-onomatopoeic word pairs for each of the 13 adjectives (Table 1, blue lines for congruent, red lines for incongruent pairs).

#### 2.3.3 Procedure

We asked participants to rate material video-onomatopoeic word pairs for the same set of adjectives. For a total of 52 types of stimuli (13 adjectives x 2 previously received word ratings; high or low x 2 previously received material ratings; high or low), we collected 13 adjective ratings, resulting in 676 total trials. All participants took part in the experiment in a sound-isolated lab. The 5-second audio recordings of the onomatopoeic words were synchronously presented in a loop with the material videos using Sennheiser SK-507364 HD 206 headphones.

The ratings were collected on a 7-point Likert scale (1 = "Not at all," 7 = "Very") with a standard cable mouse. The experiment was coded in MATLAB R2020b using Psychtoolbox and conducted on an ASUS N550J Notebook. Participants were instructed to rate each pair of onomatopoeic words and material videos based on how well they thought the word-video pair matched the adjective displayed on the screen (Figure 2).

**Figure 2.**
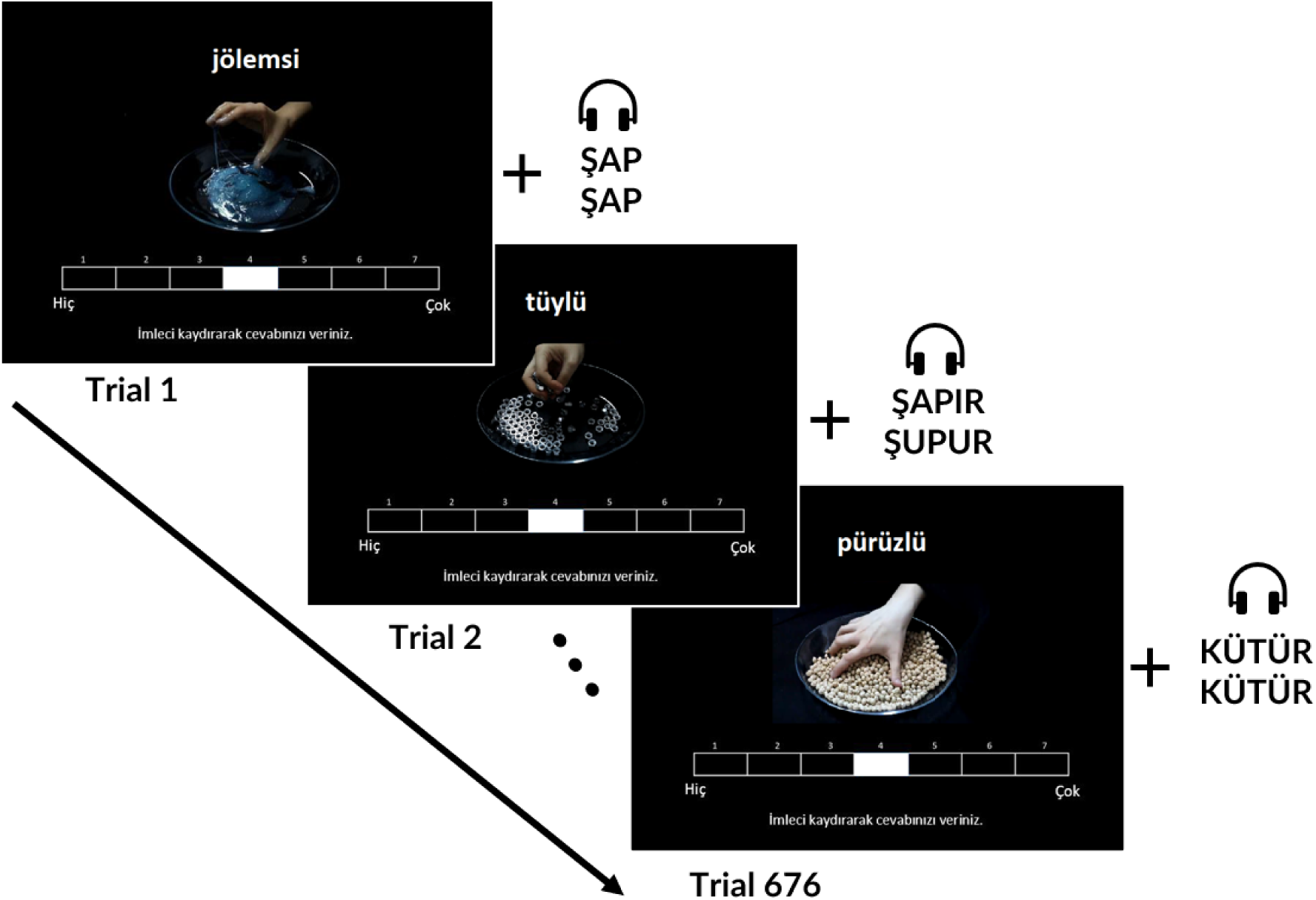
Procedure and Timeline of Experiment 2. In each trial, participants watch a 5- second material video in a loop and also listen to the audio of the Turkish onomatopoeic words in a loop while they are rating the materials.

#### 2.3.4 Results

All data analyses were performed using JASP (JASP Team, 2022). For analytical convenience, we grouped 26 individual material videos into one of the four softness dimensions (viscosity, surface softness, granularity, and roughness) and conducted the analyses at the dimension level rather than at the individual material level. A similar approach was adopted for the adjectives as well: viscosity (gelatinous, slimy, sticky, gooey, slippery, moisturous), surface softness (silky, velvety, hairy), granularity (sandy, powdery, granular), and roughness (roughened) adjectives were grouped into their corresponding softness dimension. Analyses for individual adjectives were also conducted and presented in Experiment 3 Supplementary Results.

We conducted 4 Repeated Measures ANOVAs for the adjective groups for each softness dimension. Since the groups violated the normality and sphericity assumptions, we report Greenhouse-Geisser corrected values. We also used Bonferroni correction for multiple comparisons and tested significance at **α** = .0018 level.

For the viscosity adjectives, main effects for the material dimension (*F*(1.86, 53.96) = 34.42, *p* < .001, η²p = .54), and onomatopoeic word ratings (*F*(1, 29) = 25.31, *p* < .001, η²p = .47) were significant. We also observed a significant interaction between the material dimensions and the onomatopoeic word ratings (*F*(1.96, 56.88) = 35.88, *p* < .001, η²p = .55). A post hoc analysis revealed that the mean ratings of viscosity adjectives were significantly lower for viscosity materials when they are paired with low-rated onomatopoeic words (*M* = 2.38, *SD* = 0.9) compared to when they are paired with high-rated onomatopoeic words (*M* = 3.96, *S* = 0.73; Figure 3A).

**Figure 3.**
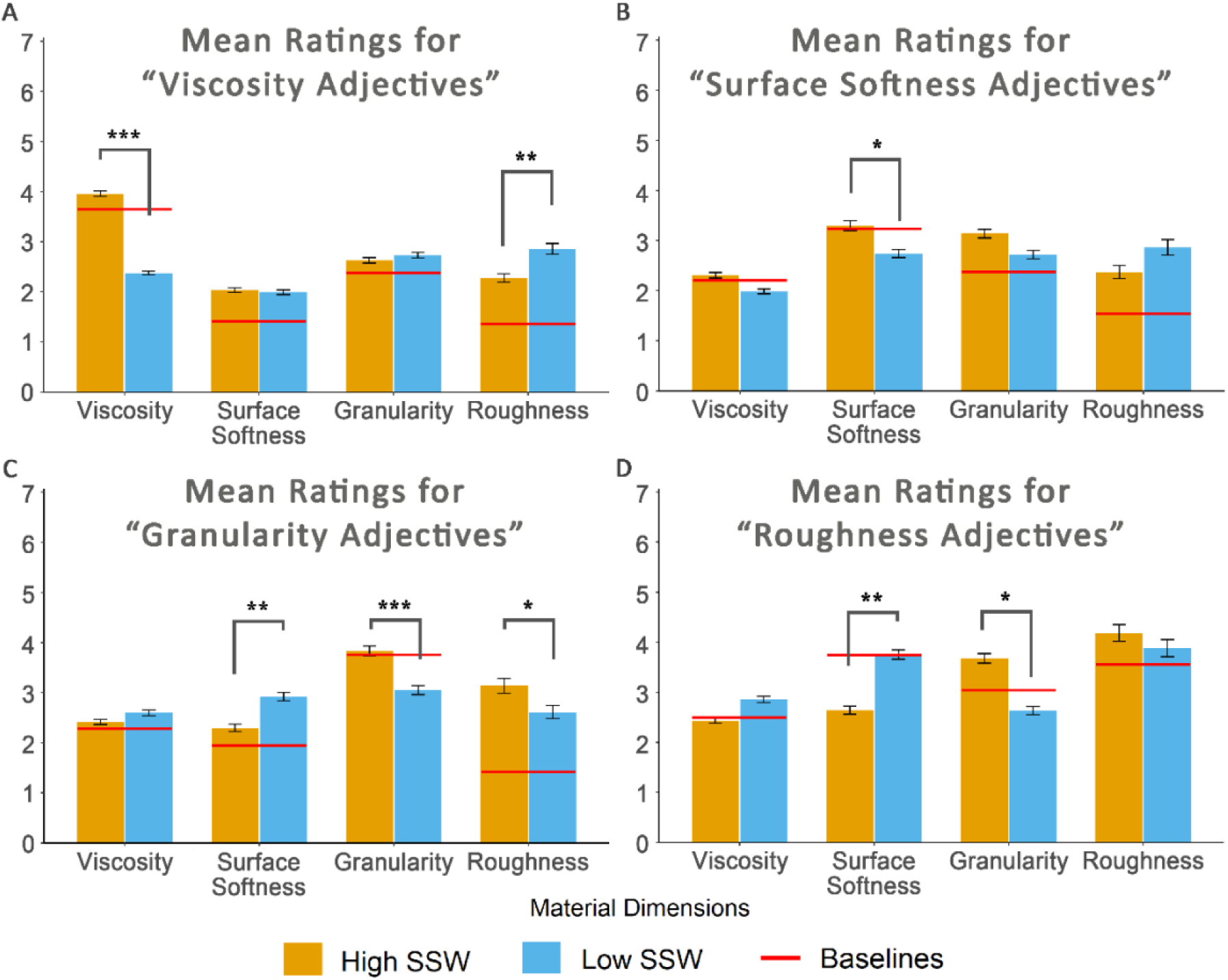
Ratings of Adjective Dimensions Across Different Cases. Yellow bars denote the conditions with high-rated onomatopoeic words while blue bars indicate the ones with low-rated onomatopoeic words. Red lines are baseline mean ratings for the corresponding material dimensions. * p < .05 ** p < .01 *** p < .001

For instance, when the Viscosity Materials such as shower gel, wool, slime, and matchstick (Table 1) are presented with their respective low-rated onomatopoeic words (e.g., “tak tak, tangır tungur, or tıkır tıkır”), they are rated lower on the viscosity adjectives (e.g., gelatinous, slimy, gooey, slippery) compared to when they are presented with their respective high-rated onomatopoeic words (e.g., “şap şap, şapır şupur, or şıpır şıpır”). On the other hand, the opposite effect was true for the roughness materials: We observed significantly higher mean ratings for the Viscosity Adjectives when they were paired with low-rated onomatopoeic words (*M* = 2.86, *SD* = 1.3) compared to when they were paired with high-rated onomatopoeic words (*M* = 2.28, *SD* = 1.11).

The surface softness adjectives also exhibited significant main effects for the material dimensions (*F*(2.61, 75.66) = 19.72, *p* < .001, η²p = .41), and onomatopoeic word ratings (*F*(1, 29) = 6.61, *p* = .016, η²p = .19). In addition, a significant interaction effect was observed with these two factors (*F*(2.19, 63.61) = 9.01, *p* < .001, η²p = .24; Figure 3B). Post hoc analyses for surface softness adjectives revealed that mean ratings were significantly lower for the surface softness materials when they were paired with low-rated onomatopoeic words (*M* = 2.74, *SD* = 1) compared to when they were paired with high-rated onomatopoeic words (*M* = 3.3, *SD* = 1.1). However, this result was carried mainly by the adjective "silky" since it was the only one with a significant difference in mean ratings between high-and low-rated onomatopoeic words (see Figure Supplementary Figure 5G).

In granularity adjectives, both main effects for the material dimensions (*F*(2.38, 69.02) = 22.02, *p* < .001, η²p = .43) and onomatopoeic word ratings (*F*(1, 29) = 8.35, *p* = .007, η²p = .22) showed significant differences. A significant interaction effect was also observed between material dimensions and word ratings (*F*(1.77, 51.25) = 14.82, *p* < .001, η²p = .34; Figure 3C). The post hoc analyses using Bonferroni correction revealed three groups with significantly different means for granularity adjectives. Firstly, similar to the Viscosity Adjectives and Surface Softness Adjectives, the mean ratings of Granularity Adjectives were significantly lower for granularity materials when they were paired with low-rated onomatopoeic words (*M* = 3.05, *SD* = 1.02) compared to when they are paired with high-rated onomatopoeic words (*M* = 3.83, *S* = 0.92). This effect is already evident in individual granularity adjectives (see Supplementary Figure 5J-L). Secondly, the mean ratings of Granularity Adjectives were significantly higher for the Surface Softness materials when paired with low-rated onomatopoeic words (*M* = 2.92, *SD* = 1.05) compared to when they were paired with high-rated onomatopoeic words (*M* = 2.29, *SD* = 1.01). This trend can be seen in the individual granularity adjectives in Supplementary Figure 5J-L, with only the adjective “granular” having a significant difference in the corresponding mean ratings. Lastly, the mean ratings of granularity adjectives were significantly lower for roughness materials paired with low-rated onomatopoeic words (*M* = 2.61, *SD* = 1.26) as opposed to when they are paired with high-rated onomatopoeic words (*M* = 3.14, *SD* = 1.36). Despite this difference, none of the individual granularity adjectives showed significant differences in ratings for roughness materials.

Finally, for the roughness adjective, significant main effects were observed for the material dimensions (*F*(2.8, 81.2) = 25.58, *p* < .001, η²p = .47) and onomatopoeic word ratings (*F*(1, 29) = 0.2, *p* = .656, η²p = 0). Additionally, a significant interaction effect was observed between these two factors (*F*(1.64, 47.58) = 8.72, *p* = .001, η²p = .23; Figure 3D). The post hoc analysis results revealed that the mean ratings were significantly higher for surface softness materials when combined with low-rated onomatopoeic words (*M* = 3.76, *SD* = 1.47) compared to when they are combined with high-rated onomatopoeic words (*M* = 2.64, *S* = 1.24). In addition, mean ratings were significantly lower for granularity materials when they were presented with low-rated onomatopoeic words (*M* = 2.63, *SD* = 1.17) compared to when they were presented with high-rated onomatopoeic words (*M* = 3.68, *SD* = 1.14).

## 3 Discussion

The non-arbitrary nature of language is evidenced by associations of certain phonemes with different shapes and sizes across cultures. More recent findings even suggest nuances in phoneme-surface quality relationships in Japanese onomatopoeia (Sakamoto & Watanabe, 2018; Wong et al., 2022; Hanada, 2023). Here, we show for the first time that Turkish onomatopoeic words have unique associations with material softness qualities. Besides, the sound symbolism effect goes beyond surface material qualities to the perception of three-dimensional everyday materials. Finally, we demonstrate that spoken onomatopoeic words can be used to manipulate participants’ softness perception of everyday materials in a dimension-specific fashion. In two preliminary studies, we examined semantic spaces of Turkish onomatopoeic words and material videos with regard to the softness properties of materials. From these results, we created congruent and incongruent word-video pairs. Next, in two experiments, we used onomatopoeic words to selectively alter adjective ratings for materials.

We observed increased ratings for the dimension-related adjectives of the congruent pairs with high-rated sound symbolic words and the opposite for incongruent pairs with low-rated sound symbolic words.

Cross-modal interactions between language and sensory processes provide some of the most striking examples of top-down influences on perception. One of the popular research topics on the subject, sound symbolism, has often focused on interactions between phonetic characteristics of words and shape perception (e.g., Bouba/Kiki effect). Only a few recent studies show a relationship between sound symbolic words and tactile material characteristics, and these are strictly conducted in Japanese (Watanabe et al., 2012; Lo et al., 2017). Similar to the haptic perception of materials, these studies also mostly focus on the surface properties of materials, with a couple of three-dimensional exceptions, such as compliant stimuli, i.e., springs (Sakamoto & Watanabe, 2017). Here, we add to surface softness properties and provide novel evidence for the overlap between multiple dimensions of perceived softness qualities of everyday materials and from Turkish sound symbolic words (SSWs). We extract four softness dimensions common to both materials and SSWs: Viscosity, surface softness, granularity, and roughness. This finding gives us the first insight into the multiple dimensions of softness via SSWs. One of the discrepancies with the literature is the lack of a deformability dimension for SSWs in our findings. Unlike material videos where we include deformable materials such as playdough, the word list in Supplementary Study 2 does not include any SSWs that correspond to the sound of a deforming material. To our knowledge, Turkish has no such examples, most likely because deforming materials do not make any sound (e.g. when compared to splashing water).

Next, we use these Turkish SSWs to manipulate the perceived softness qualities of materials along multiple softness dimensions. By pairing high-and low-rated onomatopoeic words with various materials, we observe changes in the ratings of adjectives related to the materials’ softness dimensions. Compared to baseline ratings (video-only stimuli), these dimension-specific changes are in the same direction as the ratings of the onomatopoeic words.

Our findings reveal fluctuations in dimension-specific adjective ratings predicted by the ratings for SSWs. Pairing viscosity materials with low-rated onomatopoeic words results in significantly lower ratings of viscosity-related adjectives (but not for adjectives in other dimensions) compared to when they were paired with high-rated words. Here, we also see a surprising effect, where the SSWs change the roughness materials’ ratings of slimy and slippery adjectives in an unexpected direction (Supplementary Figure 5B and E). Participants’ slimy and slippery ratings for “sandpaper” are higher when presented with a low-rated SSW for ‘roughened’ (gıcır gıcır) compared to when presented with a high-rated SSW for ‘roughened’ (kırt kırt). Except for ‘moisturous,’ all other viscosity adjectives follow this trend as well, supporting the negative correlation between roughness and viscosity dimensions. An important observation for the viscosity adjectives dimension was that the effect of SSWs on the adjective ratings appeared to have a stronger diminishing impact than an enhancing one, which requires further research to understand the potential asymmetry of sound symbolism’s cross-modal effects.

For the surface softness dimension, SSW-material pairings only affected the ratings for surface softness materials significantly, as expected (Figure 3B). This result seems to be carried by the adjective silky: the ratings for velvety and hairy also differed in the expected direction, but these differences did not reach a significance level (Supplementary Figure 5G, H, and I).

The SSWs altered all the granularity adjective ratings (sandy, powdery, and granular) in the expected direction for the granularity materials. The hairy ratings for granularity materials (Supplementary Figure 5I) were also affected similarly. Even though hairy is an adjective describing surface softness characteristics, it might indirectly be influenced by an SSW meaning, e.g., “püfür püfür” describing the gentle and refreshing wind that is blowing softly and coolly or “pıtır pıtır” (pitter patter) might have caused the granularity materials to be associated with the hairiness. Another unforeseen difference was the surface softness materials’ ratings for the granularity adjectives. Here, incongruent SSWs such as katır kutur or kıtır kıtır might mean a crunchy sound as in apple or a crispy sound as in cornflakes. Having this in mind, it is not surprising that for some participants, these SSWs resulted in higher semantic associations with being granular.

Finally, for the roughness adjective, both granularity and roughness materials showed effects in the expected direction, but the difference for roughness materials remained insignificant. Each dimension had different numbers of stimuli to be rated by the participants since our selection of adjectives, materials, and onomatopoeic words was based on the PCA results. This was especially true for roughness. As a result of the preliminary studies, we were able to choose only one adjective (roughened) with two corresponding material videos and two onomatopoeic words, compared to three to six adjectives in other dimensions. Thus, the insufficient number of ratings might have caused these results for the roughness adjective. This dimension is considered to be a control since it describes the surface qualities and not three-dimensional softness characteristics such as deformability (Dövencioğlu et al., 2022). So, our starting point to investigate sound symbolism for material qualities beyond surfaces might have led us to bring less focus to this dimension. Still, future research should aim to have a balanced number of stimuli across dimensions, if at all possible.

We also observed a significant difference in the mean ratings of the roughness adjective for the surface softness materials in the opposite direction to the roughness materials when paired with high and low-rated onomatopoeic words. This means that onomatopoeic words rated high in surface softness dimension decreased the roughness ratings of the surface softness materials - which is not surprising. This finding shows a clear contrast of the effects of sound symbolic words between the surface softness and roughness dimensions.

Another curious contrast we observed was in both the ratings for viscosity and surface softness adjectives. In both cases, we observed opposite effects for roughness materials, i.e., pairs with high-rated SSWs had lower rating values compared to the pairs with low-rated SSWs. This might be because a high-rated roughness material (sandpaper) paired with a low-rated SSW (gıcır gıcır meaning crisp, shiny, or brand new) might result in a higher rating for surface softness properties such as silky. It might also mean that the semantic association of the SSW and the material is conflicting, that the participants rated only one, either the SSW or the video. This distinction cannot be made conclusively from this study and needs further testing. Nevertheless, both types of contrasts suggest that a single SSW can sometimes provide information for more than one softness dimension, and sound symbolic effects on material perception are more complex than initially thought.

Lastly, our findings suggest an interesting parallel between the effects of sound symbolism on granularity and roughness adjectives when paired with their corresponding materials. This observation suggests a potential overlap between the two softness dimensions and brings up a new line of questioning for future research.

Overall, our results support the hypotheses that within-dimension pairs, matching materials with high-rated onomatopoeic words would enhance the mean ratings of adjectives related to the materials’ softness dimensions. In contrast, when materials are paired with low-rated onomatopoeic words, the mean ratings of the adjectives that describe the materials’ softness dimensions will decrease. Collectively, these findings deepen our understanding of the complex interplay between sound symbolism and material perception, particularly for perceived softness. They present promising pathways for further research while highlighting the unique role of onomatopoeic words in material perception. This area of research may benefit from cross-cultural studies exploring both the phonetic features of onomatopoeic words and material perception. The universality of the Bouba/Kiki effect suggests that these phonetic effects might have commonalities in different cultures. The question of whether these findings are limited to Turkish speakers is a potential start for following research future investigations.

## 4. Acknowledgments

This study is supported by ODTÜ BAP (AGEP-104-2022-10910) and TÜBİTAK 1001 (122K914) projects awarded to DND.

## 1 Supplementary Study 1

### 1.1 Participants

15 university students (*M* = 23.8, *SD* = 4.5, 2 Males) volunteered to participate in the online study (Qualtrics) through the Middle East Technical University research sign-up system. All participants were naive to the purposes of the study. Only one participant was left-handed. Participants gave written informed consent before the study and were debriefed and given course credit after they had completed the study.

### 1.2 Stimuli

#### 1.2.1 Material Videos

We used material videos as stimuli as they are reported to be assessed as comparable to haptic stimulus judgments (Baumgartner et al., 2013; Cavdan et al., 2021; Okamoto et al., 2013). Thirty-two soft and 8 hard materials (as control) were selected (see Supplementary Figure 1 for the complete list of materials). Videos showed materials in their glass containers and experimenters’ hand movements against a black background. Videos were recorded without sound using a Canon EOS M50 camera with a tripod placed 50 cm away from the materials. Natural light and automatic ISO settings were used. The spatial resolution was set to 1920’1080 pixels Full HD, and videos were recorded at 50 frames per second. Next, 5-second video clips were cut for the best representative exploratory procedures that present the most salient feature of a material, such as pressing for sponge pieces (deformable), rubbing for velvet (textile), stirring for hand cream (viscous, Dovencioğlu et al., 2022).

**Supplementary Figure 1.**
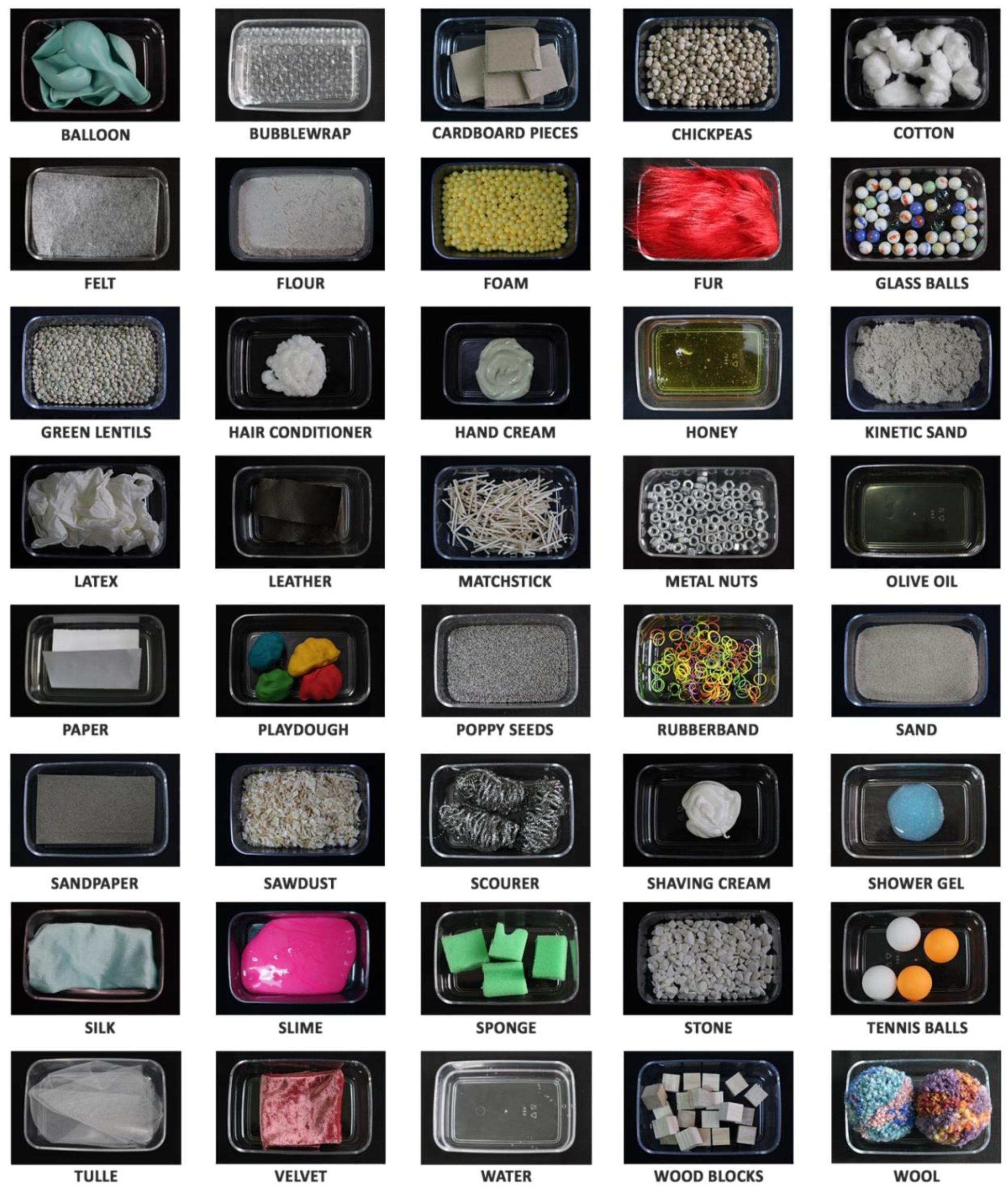
All Materials Used in Study 1.

#### 1.2.2 Adjectives

The list of 31 adjectives used here is taken from a previous study that was initially adapted to Turkish by Dövencioğlu et al. (2019; 2022) from Guest et al. (2011). This list consisted of non-sensual adjectives that relate to material characteristics of soft and rough materials. We only removed 2 adjectives from the list, “meaty” and “leathery." The former was not included since it was not relevant to our material list, and the latter was not included since the Turkish translation of the adjective was not a commonly used word in the daily language. The complete list of the resulting 29 adjectives and their English translations can be found in Supplementary Table 1.

**Supplementary Table 1.**
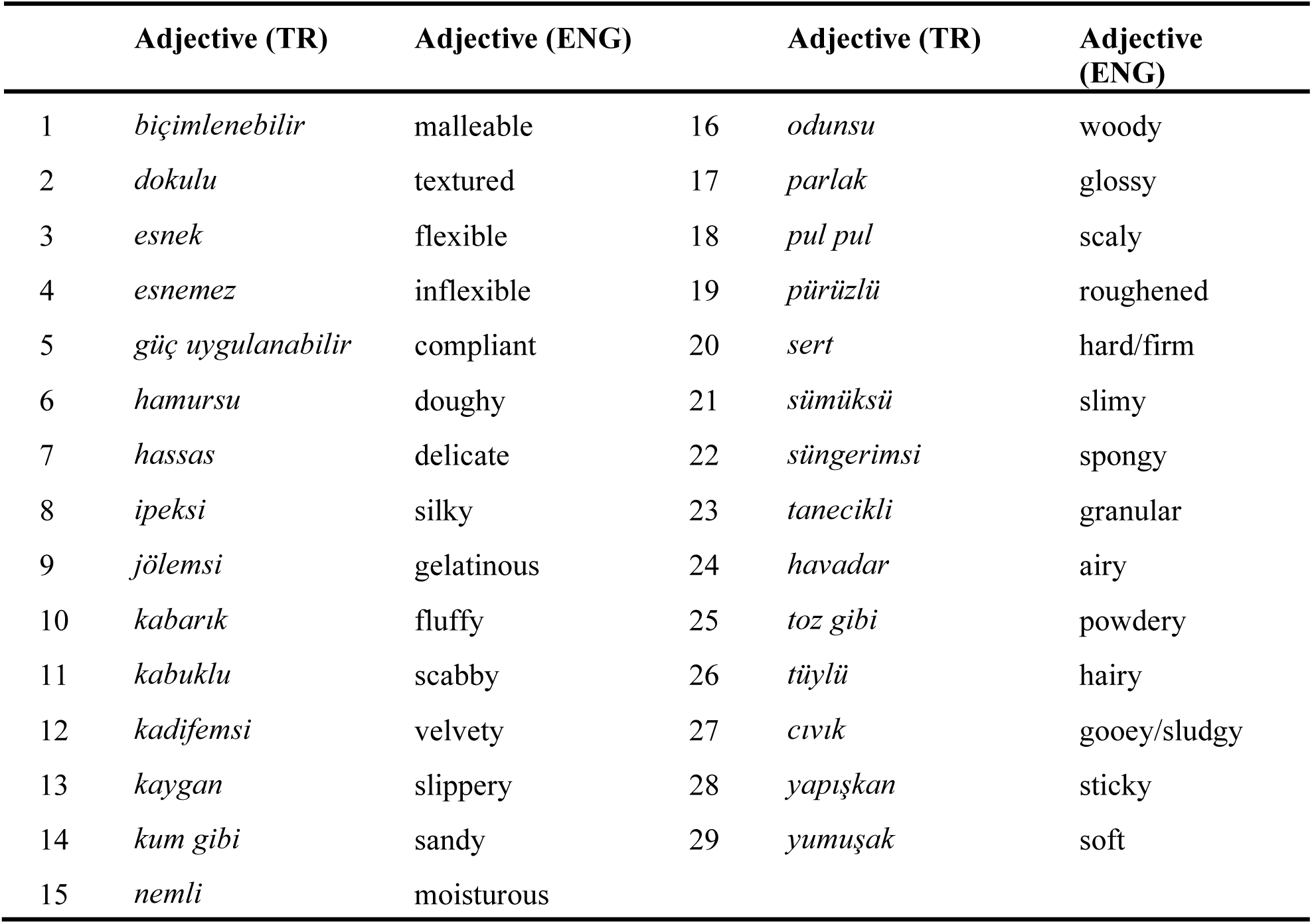
29 Adjectives Used in Study 1 with Their English Translations.

### 1.3 Procedure

The experiment was conducted online on Qualtrics due to the global COVID-19 pandemic restrictions. Participants were instructed to play the 5-second video of the material and then to rate the materials shown in the videos based on the adjectives. Overall, they completed 40 blocks with 29 adjectives each for one material video. They were required to complete all ratings before proceeding to the next trial. Adjectives were rated on a 7-point Likert scale (1 = "*Not at all*”, 7 = “*Very appropriate*”). Supplementary Figure 2 depicts a sample trial from the study.

**Supplementary Figure 2.**
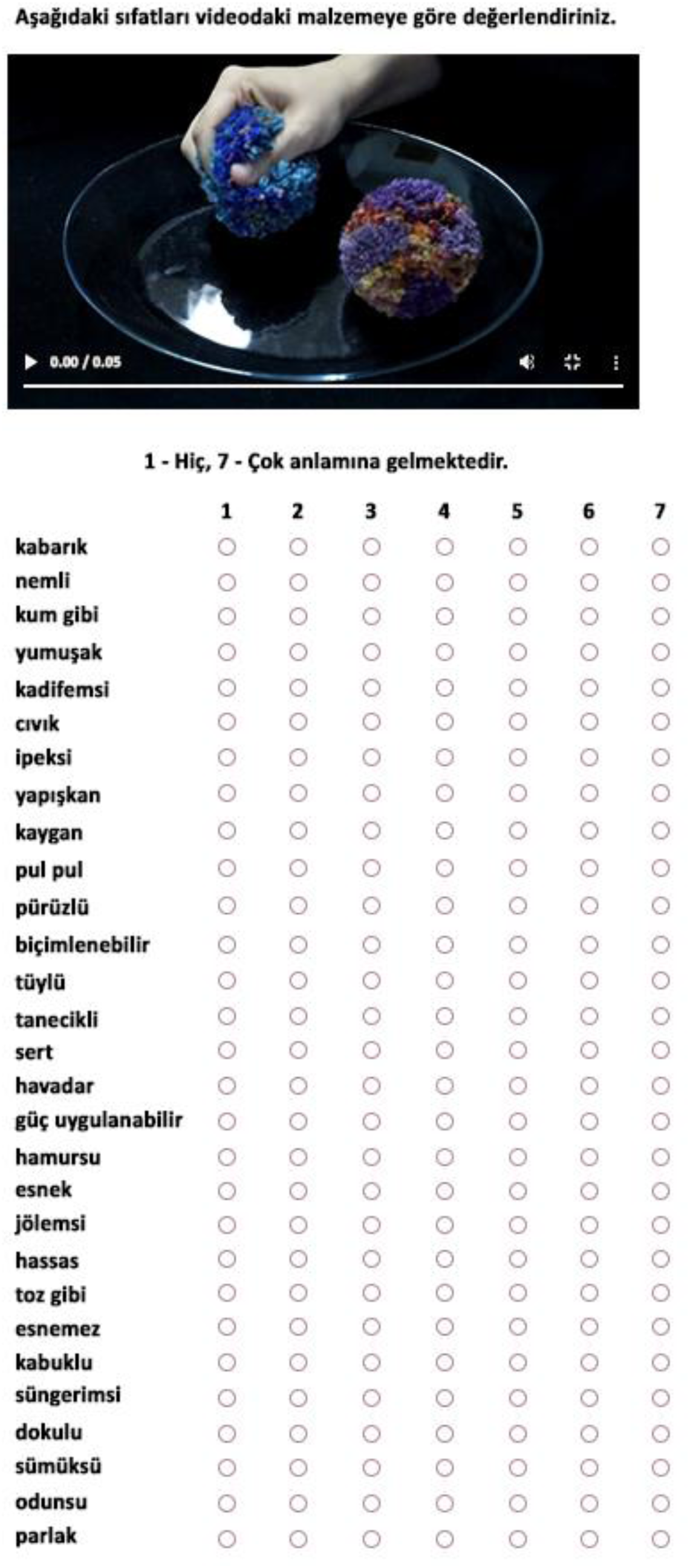
Sample Single Trial from the Study 1.

### 1.4 Results

Before the analysis, we ran Barlett’s test of sphericity (χ2(406) = 1546, *p* < .001) and Kaiser-Mayer-Olkin (KMO) test (score = .584) to make sure that the data can be used in a Principal Component Analysis (PCA). The PCA was conducted to see how adjectives load to each softness dimension using JAMOVI (The Jamovi project, 2019). Using the Kaiser normalization and varimax rotation, 7 principal components were extracted, explaining 87.58% of the total variance in the data. The component loadings are shown in Supplementary Table 2.

**Supplementary Table 2.**
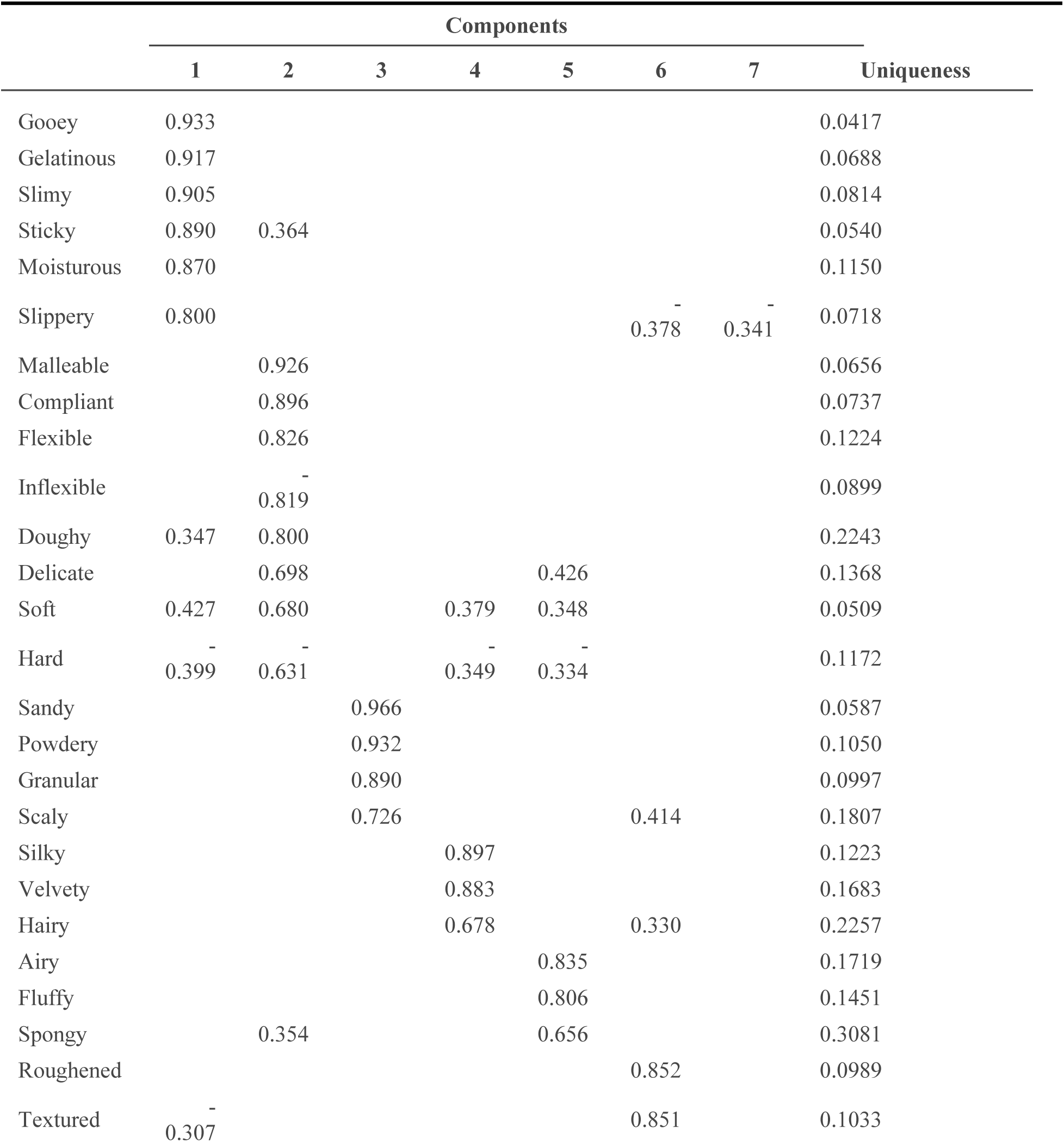
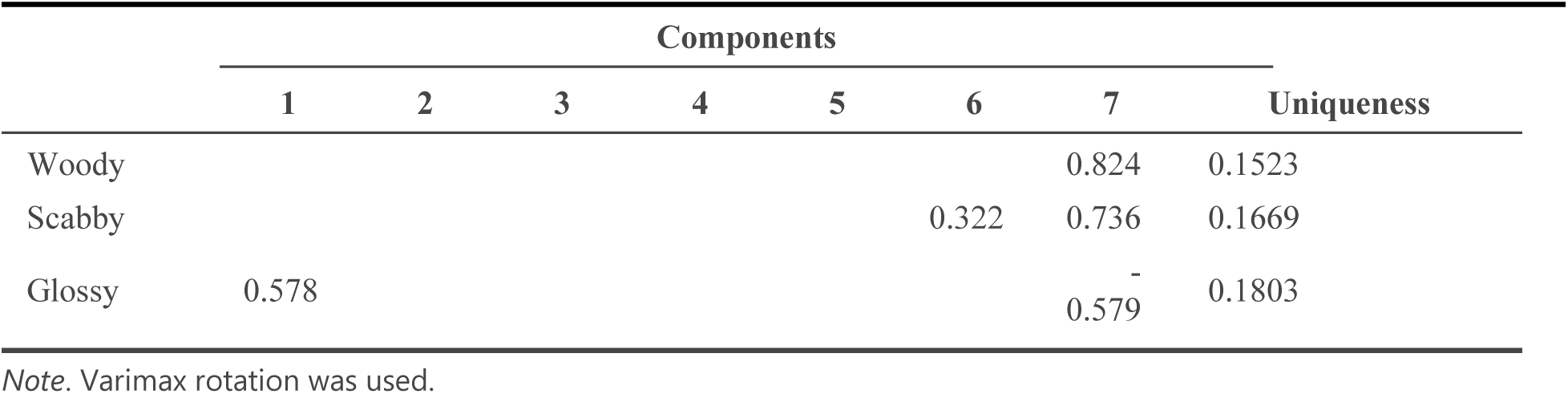
PCA Results for Study 1.

The first component, Viscosity, accounted for 21.03% of the variance and was characterized by the adjectives *gooey, gelatinous, slimy, sticky, moisturous, slippery,* and *glossy.* The second component, Deformability, accounted for 20.32% of the variance and comprised the adjectives *malleable, compliant, flexible, inflexible (-), doughy, delicate, soft,* and *hard (-)*. The third component, Granularity, explained 12.43% of the variance and included the adjectives *sandy, powdery, granular*, and *scaly*. The fourth component, Surface Softness, accounted for 9.17% of the variance and included the adjectives *silky*, *velvety*, and *hairy*. The fifth component, Fluffiness, explained 8.93% of the variance and consisted of the adjectives *airy, fluffy,* and *spongy*. Roughness, the sixth component, explained 8.4% of the variance and included the adjectives *roughened* and *textured*. Lastly, the seventh component, Scabbiness, accounted for 7.3% of the variance and was characterized by the adjectives *woody* and *scabby*.

## 2 Supplementary Study 2

### 2.1 Participants

30 university students (*M* = 22.4, *SD* = 2.4, 7 males) volunteered through the Middle East Technical University research sign-up system. Three participants were left-handed, and all participants had normal or corrected to normal vision. None of the participants reported a hearing loss or any other auditory condition. All participants were native Turkish speakers and naive to the purposes of the study. An informed consent was provided before the study, and participants received course credits after completion.

### 2.2 Stimuli

#### 2.2.1 Spoken Onomatopoeic Words

First, 51 material-related onomatopoeic words were chosen from Turkish linguistics literature (Zulfikar, 1995; Ozkan, 2010). After that, we checked the meanings of these words from the Turkish Language Society, TDK (Türk Dil Kurumu) dictionary. We also checked the frequencies of the words using TS Corpus, the largest Turkish corpora (Sezer, 2017; Sezer & Sezer, 2013). Those with a frequency of 0 were eliminated since they might be unfamiliar to the participants. We completed another online rating study to determine the familiarity of the words and further eliminated the words with an average familiarity rating below 3 (out of 7). In the last step, we removed the words that are not related to our material list (e.g., “vız vız” referring to the sounds of the bees), resulting in a list of 27 onomatopoeic words (see Supplementary Table 3 for the complete list of onomatopoeic words).

**Supplementary Table 3.** Onomatopoeic Words Used in Study 2.

|  | **Adjective (TR)** | **Explanation (ENG)** |
| --- | --- | --- |
| 1 | çıt çıt | Snap fastener, gripper |
| 2 | efil efil | Gently, intermittently, and slowly (blowing wind, snowing, hair waving) |
| 3 | gıcır gıcır | Crisp, brand new |
| 4 | haşır huşur | Hard and dry things wring, wheezing, rumbling |
| 5 | hışır hışır | With a rustling sound |
| 6 | katır kutur | With a harsh and rough sound. |
| 7 | kırt kırt | Light, crisp sound, small or delicate breaking or cracking |
| 8 | kıtır kıtır | Crispy, brittle, crusty |
| 9 | kütür kütür | Crisp, fresh, with a crunching sound |
| 10 | lıkır lıkır | With a gurgling sound |
| 11 | lime lime | In small pieces, rags, and tatters |
| 12 | mırıl mırıl | Murmuring |
| 13 | mışıl mışıl | Sleeping peacefully and soundly, with a quiet and deep breath |
| 14 | pıtır pıtır | With a patter |
| 15 | pofur pofur | Soft, muffled, and continuous sound |
| 16 | püfür püfür | Light, gentle breeze or the sound of air softly moving or rustling |
| 17 | şap şap | Kissing with a screed sound, alum alum |
| 18 | şapır şupur | The sound of "smack-whisk" when kissing or eating |
| 19 | şarıl şarıl | Flowing splashingly, with a splashing sound |
| 20 | şıp şıp | Making a 'flashing' sound, plop |
| 21 | şıpır şıpır | Water droplets falling; a gentle, rhythmic dripping sound |
| 22 | şırıl şırıl | Continuous and loud flowing of water with a pleasant noise |
| 23 | tak tak | The sound that is made during hitting, impact, rat-tat |
| 24 | tangur tungur | Crash bang wallop, bone-shaking, clack |
| 25 | tıkır tıkır | At a rattling pace, tickety-boo |
| 26 | tiril tiril | Crisp and clean, gauzy, floaty |
| 27 | vıcık vıcık | Ropy, sludgy, gooey, slushy |

One of the experimenters (BMH, a native Turkish speaker) recorded the spoken onomatopoeic words using a RODE noise-canceling microphone. The recordings took place in a soundproof room and were edited using Audacity software (Audacity Team, 2021). The audio files started with a 1-second silent onset before the onomatopoeic words (∼2 seconds), followed by a silent offset time to complete the recordings to a duration of 5 seconds. The audio recordings were normalized for volume level consistency in the final step.

#### 2.2.2 Adjectives

The adjective list used in Study 1 is shortened in Study 2 to optimize experiment runtime. We removed the deformability related adjectives (malleable, compliant, flexible, inflexible, doughy, delicate, textured, spongy) in Study 2.

### 2.3 Procedure

The study was coded in MATLAB R2020b (The MathWorks Inc., 2020) using Psychtoolbox-3 and included 27 spoken onomatopoeic words along with 21 adjectives. Participants rated every onomatopoeic word for 21 adjectives, resulting in a total of 567 randomized trials for a single experimental session. The experiment was conducted using an HP ENVY dv6 Notebook in a sound-isolated room. The 5-second audio recordings of the onomatopoeic words were presented to the participants synchronously with their written forms on the screen in a loop, using Sennheiser SK-507364 HD 206 headphones. Ratings were collected with a standard cable mouse after the first 3 seconds of the recordings to ensure that the participants listened to the recordings. A 7-point Likert scale was used for the ratings (1 = "*Not at all”*, 7 = “Very”). All participants were instructed to listen to the audio first and then rate the onomatopoeic word for the adjective they saw written on the screen. A sample trial is depicted in Supplementary Figure 3.

**Supplementary Figure 3.**
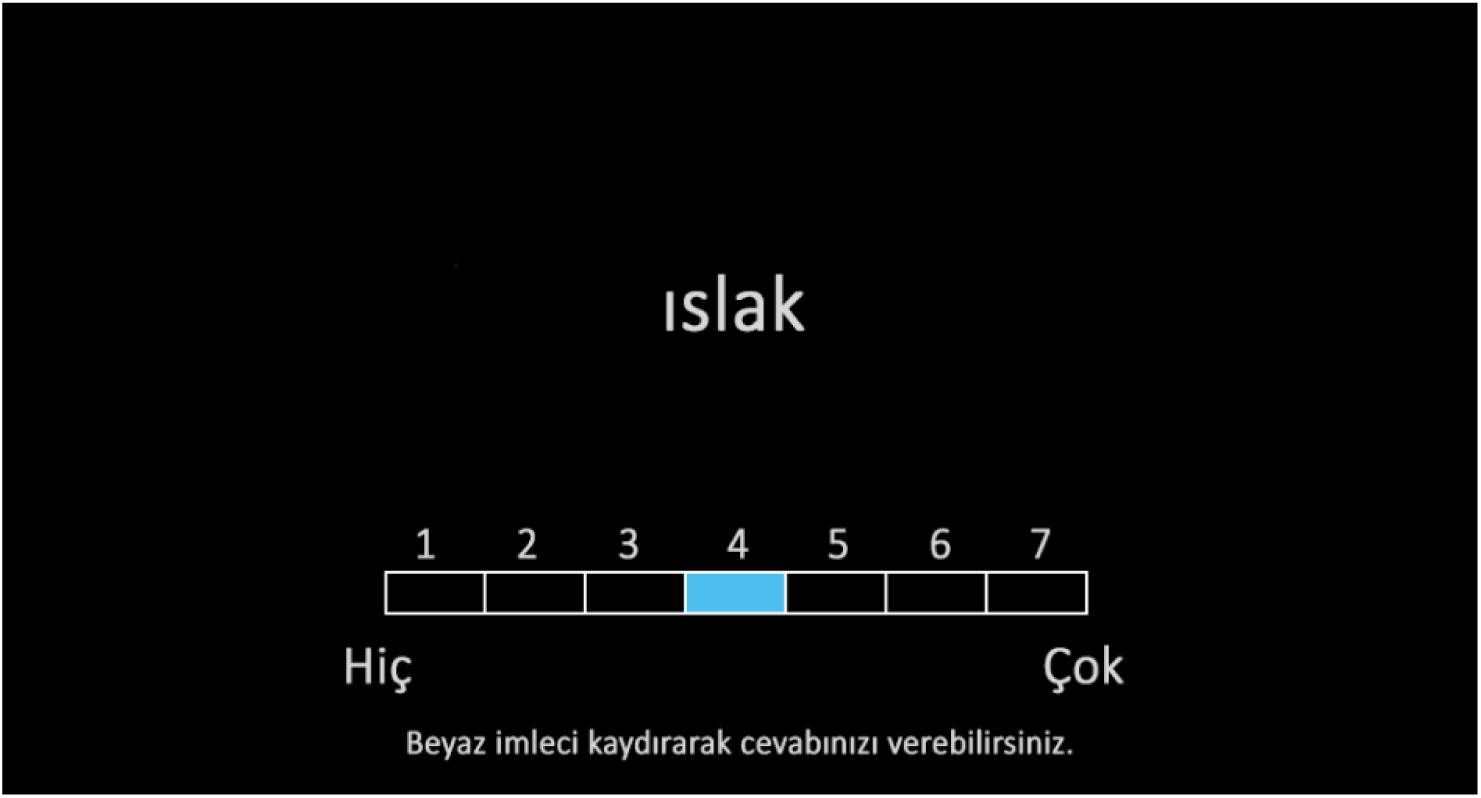
Sample Trial from the Study 2. The adjective “wet” is presented on the screen while the participant is hearing the onomatopoeic word from the headphones. A rating scale is presented to rate the adjectives using the cursor.

### 2.4 Results

A Principal Component Analysis was conducted to see the resulting adjective dimensions using JAMOVI (The Jamovi project, 2019). Barlett’s test of sphericity was significant (χ2(210) = 995, *p* < .001), and the Kaiser-Mayer-Olkin (KMO) test resulted in a score of .535, indicating that the data is suitable for a PCA. Using the Kaiser normalization and varimax rotation, 4 principal components were extracted. The resulting component loadings are presented in Supplementary Table 4.

**Supplementary Table 4.**
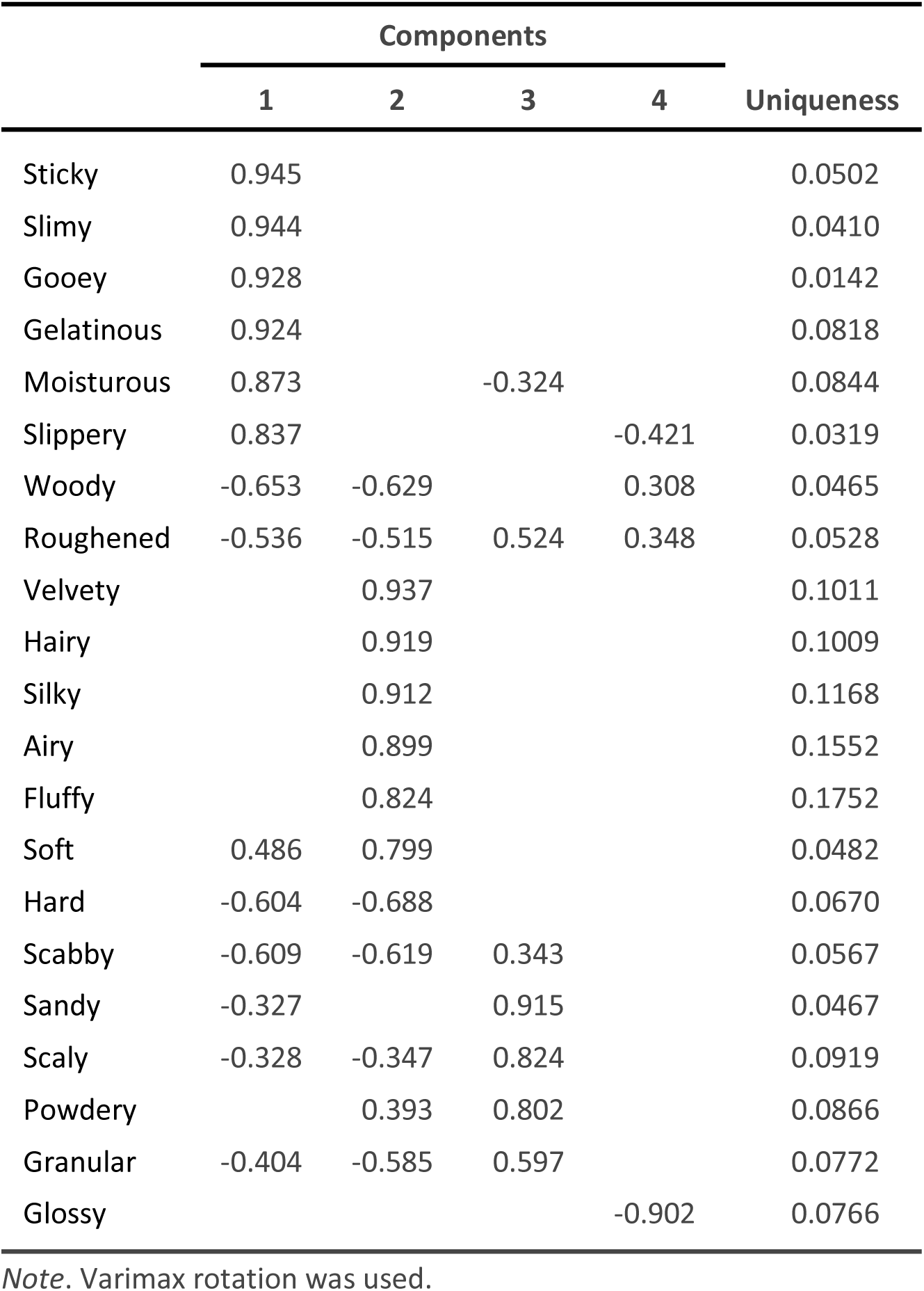
PCA Results for the Study 2.

The first component, Viscosity, accounted for 35.02% of the variance, and it was characterized by the adjectives *sticky, slimy, gooey, gelatinous, moisturous,* and *slippery.* The second component, Surface Softness, accounted for 32.91% of the variance and comprised the adjectives *velvety, hairy, silky, airy, fluffy, soft, hard (-), and scabby (-)*. The third component, Granularity, explained 16.85% of the variance and included the adjectives *sandy, scaly, powdery*, and *granular*. The fourth component, Roughness, explained 7.85% of the variance and included the adjectives *woody, roughened,* and *glossy (-)*.

## 3 Experiment 2: Supplementary Results for Individual Adjectives

First, we analyzed the effects of high and low-rated onomatopoeic words on individual adjectives across 4 different material dimensions. The results of Shapiro-Wilk tests and Mauchly’s sphericity tests showed that assumptions of normality and sphericity had been violated in all groups for 13 adjectives. We conducted 13 Repeated Measures ANOVAs using Greenhouse–Geisser adjustment and Bonferroni correction for multiple comparisons (resulting in α = .05/13 = .003). All ANOVA results except for the adjective “granular” showed a significant main effect of material dimensions (see Supplementary Figure 4A for the ANOVA results of adjectives). Only the adjectives “sticky, gooey, moisturous, hairy, and powdery” showed significant main effects of onomatopoeic word ratings. All ANOVA results showed significant interactions between the material dimensions and ratings of the onomatopoeic words. To determine which specific material dimensions and onomatopoeic word ratings caused these significant interaction effects, we conducted 4 post hoc tests for each of the 13 adjectives (see Supplementary Figure 4B).

**Supplementary Figure 4.**
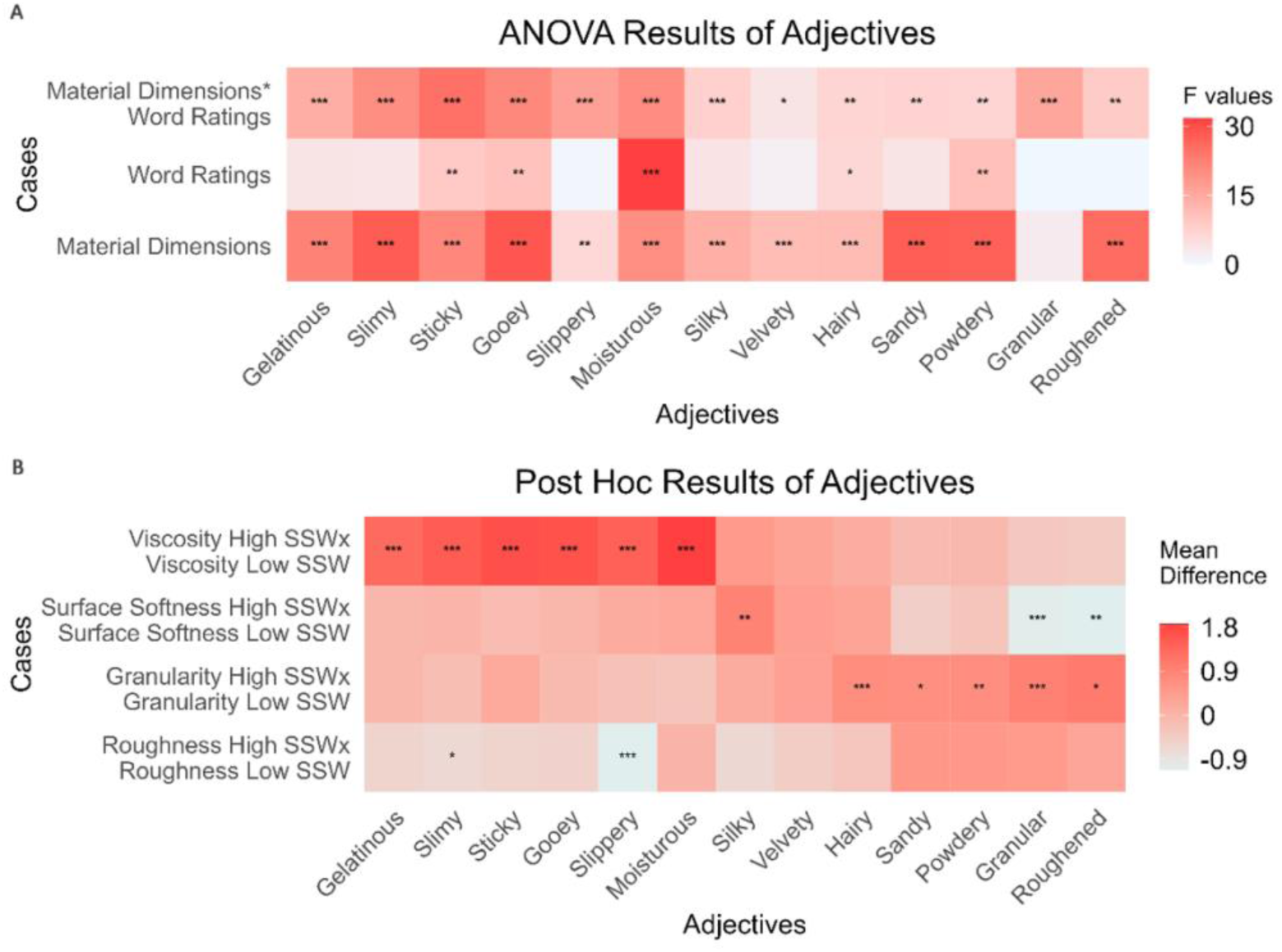
ANOVA and Post Hoc Results of Individual Adjectives. (A) Red color gradient represents the F values of ANOVAs, with white being zero and increasing towards red. Asterisks indicate the p values (* p < .05 ** p < .01 *** p < .001). (B) Red color gradient represents the mean differences in the post hoc tests, with red being a positive difference and decreasing towards -0.9 in grey. Asterisks indicate the p values (* p < .05 ** p < .01 *** p < .001).

There were significant differences between the mean ratings of all viscosity adjectives (gelatinous, slimy, sticky, gooey, slippery, and moisturous) for the viscosity materials that were matched with high or low-rated onomatopoeic words. For example, the mean ratings of the adjective "gelatinous" were lower for the viscosity materials, including "shower gel, green lentils, and hand cream" (see Table 1 for the complete list) when they were presented with their respective low-rated onomatopoeic words such as "tak tak, çıt çıt, or lime lime” as opposed to when they were presented with the high rated onomatopoeic words such as “şap şap, vıcık vıcık or şarıl şarıl” (see Supplementary Figure 5A). The adjectives “slippery” and “slimy” also had significant differences in their mean ratings for the roughness materials (Supplementary Figure 5B and 5E). More specifically, the mean ratings of "slimy" and "slippery" were significantly higher for the roughness materials (sandpaper and balloon) when they were presented with the low-rated onomatopoeic word "gıcır gıcır” compared to their mean ratings when they are presented with the high rated onomatopoeic word “kırt kırt." For the surface softness adjectives (silky, velvety, and hairy), only in "silky," there was a significant difference between the mean ratings for the surface softness materials when they were paired with high or low-rated onomatopoeic words (Supplementary Figure 5G). For the adjective "hairy," these differences in the mean ratings were observed for the granularity materials when they were paired with high or low-rated onomatopoeic words (Supplementary Figure 5I).

**Supplementary Figure 5.**
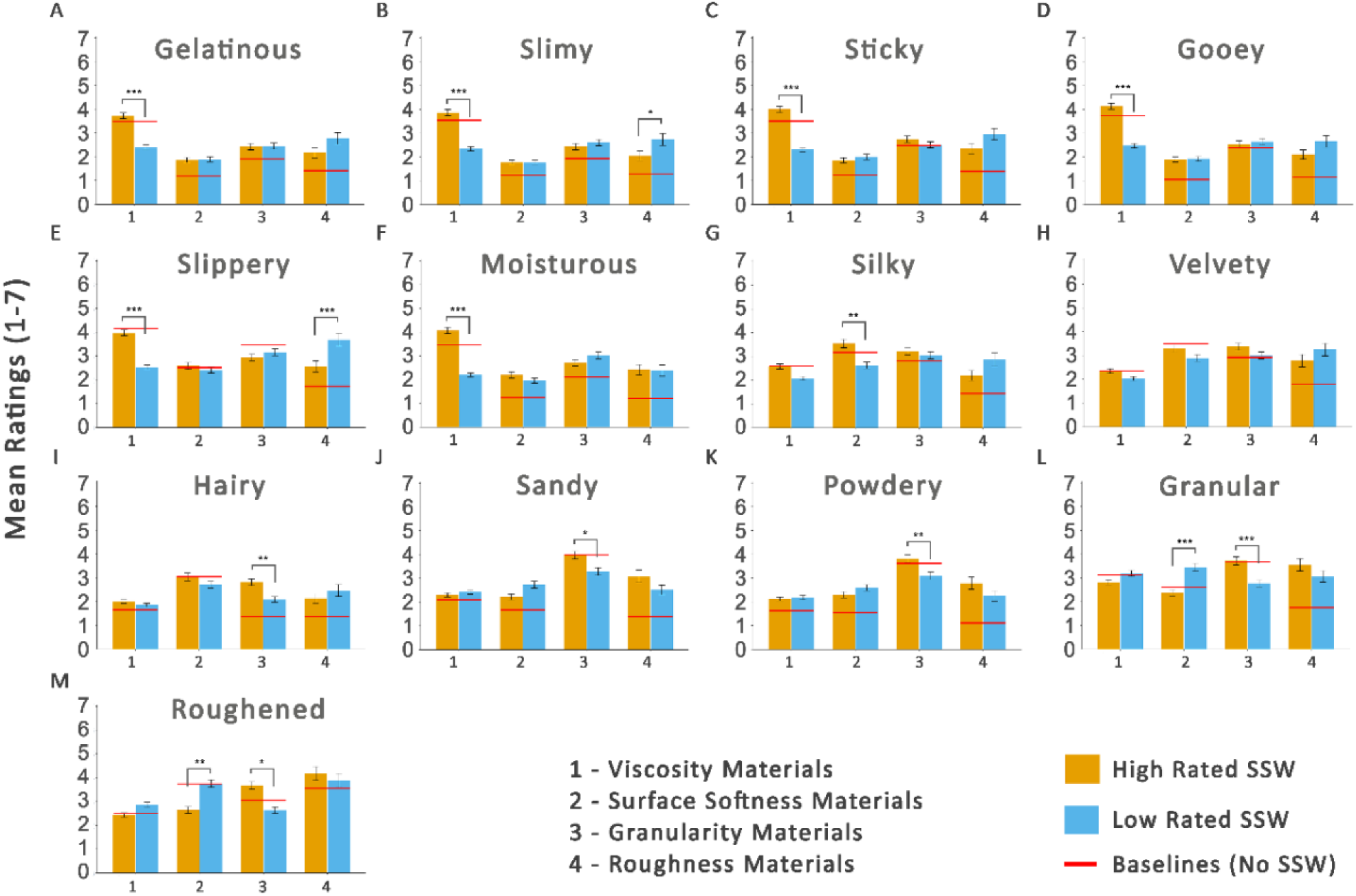
Ratings of Individual Adjectives Across Different Cases. Yellow bars denote the conditions with high-rated onomatopoeic words while blue bars indicate the ones with low-rated onomatopoeic words. Red lines are baseline mean ratings for the corresponding material dimensions. * p < .05 ** p < .01 *** p < .001

There were also significant differences between the mean ratings of all granularity adjectives (sandy, powdery, and granular) for the granularity materials that were matched with high or low-rated onomatopoeic words. For example, the mean ratings of the adjective "powdery" were lower for the granularity materials (i.e., sand, poppy seeds, latex) (Table 1) when paired with a low-rated onomatopoeic word such as “lıkır lıkır, efil efil, or şırıl şırıl” compared to when paired with a high-rated onomatopoeic word such as “hışır hışır, püfür püfür, or pıtır pıtır” (Supplementary Figure 5K). There was also a significant difference between the mean ratings of "granular" for the surface softness materials when paired with a high or low-rated onomatopoeic word (Supplementary Figure 5L).

Finally, neither low-nor high-rated onomatopoeic words affected the ratings of roughness materials for the adjective “roughened." However, there were significant differences in the ratings of surface softness and granularity materials when paired with high or low-rated onomatopoeic words (Supplementary Figure 5M). For example, the Surface Softness materials (i.e., silk, stone, or velvet) received significantly lower mean ratings for the adjective "roughened" when matched with low-rated onomatopoeic words compared to high-rated onomatopoeic words. All remaining results for the individual adjectives are illustrated in Supplementary Figure 4 and Figure 5.

